# Integrated scFv identification and CAR T cell generation for AML targeting in vivo

**DOI:** 10.1101/2024.10.08.617189

**Authors:** Yi Liu, Annika Lauk, David Sedloev, Josephine Brysting, Ela Cetin, Chunan Liu, Maximilian Mönnig, Thomas Luft, Haiyang Yun, Michael Schmitt, Tim Sauer, Fengbiao Zhou, Christian Rohde, Carsten Müller-Tidow

**Affiliations:** Department of Internal Medicine V, Hematology, Oncology and Rheumatology, Heidelberg University Hospital, 69120, Heidelberg, Germany; Molecular Medicine Partnership Unit (MMPU), European Molecular Biology Laboratory (EMBL) and Heidelberg University Hospital; Structural and Molecular Biology, Division of Biosciences, University College London, London, UK

**Keywords:** scFv, phage display library, CAR T cell therapy, targeted therapy, acute myeloid leukemia, CLL1

## Abstract

Cancer immunotherapy has witnessed remarkable advancements, especially in the development of chimeric antigen receptor (CAR) T cell therapy. Here, we integrated single-chain variable fragment (scFv) development with CAR T cell generation based on a newly developed scFv phagemid library. High-throughput long-read PacBio sequencing identified 4.5 x 10^7^ unique full-length scFv proteins within the generated library. In a proof of principle, we screened for scFvs targeting C-type lectin-like molecule-1 (CLL1) with subsequent cloning into a third generation retroviral CAR backbone. Functional assays revealed the specificity and potency of these CAR T cells in targeting CLL1-positive AML cells in vitro. *In vivo* studies reduced tumor burden and improved survival rates compared to controls. Taken together, screening for tumor specific scFvs against CLL1 can rapidly generate AML specific CAR T cells with effective tumor killing *in vivo*.

## Introduction

Antibody- and antibody fragment-mediated targeted immunotherapy is transforming cancer treatment by offering a precise and potent alternative to traditional chemotherapy^1, 2^. Single-chain variable fragments (scFvs), engineered by connecting the variable regions of the antibody’s heavy (VH) and light chains (VL) with a short peptide linker, retain the antigen-binding specificity of the original antibody and have versatile applications in cancer therapy, including monoclonal antibody development, targeted drug delivery, and CAR T cell therapy^3–8^. FDA has approved six CAR T cell therapies incorporating scFvs for the treatment of hematologic malignancies, though not for myeloid leukemia ^2,9^. The success of CAR T cells depends on the specificity and affinity of scFvs, making the development of robust scFv libraries fundamental ^10–13^. Despite several existing scFv-phage libraries have been created, some face limitations stemming from a restricted donor pool, insufficient quality assessment, or a lack of evidence supporting their therapeutic uses^14–17^.

Acute myeloid leukemia (AML) is an aggressive blood cancer characterized by the abnormal proliferation of myeloid cells. It requires prompt and effective treatments. Traditional chemotherapy remains the standard of care for many AML patients, but associated with significant toxicity and limited efficacy in certain subgroups, such as older adults or those with relapsed or refractory disease ^18–20^. In recent years, targeted therapy has emerged as a promising approach to address the challenges posed by AML^21–23^. Targets for therapeutic intervention in AML include antigens such as CD33, CD123, CD70, and CLL1^7,24–27^. CLL1 (C-type lectin-like molecule-1), also known as CLEC12A, has emerged as an attractive target because CLL1 is regularly not or very low expressed on healthy hematopoietic stem cells or other tissues but is expressed on myeloid leukemia cells, including leukemic stem cells^26–29^. Therefore, targeting CLL1 may provide a specific and selective way to treat AML.

In this study, we generated an scFv library by introducing the full repertoire of scFv genes into the genome of bacteriophages. Using third-generation PacBio sequencing technology^30–32^, known for generating long reads (>10k pb) with high fidelity and allowing the retrieval of full-length scFv (∼ 800bp), we thoroughly examined the library’s coverage and diversity. Panning efforts yielded unique scFvs with specificity for CLL1. These scFvs were then used to engineer CAR T cells, exhibiting robust *in vitro* and *in vivo* cytotoxicity against CLL1-positive tumor cells. The proven functionality of anti-CLL1 scFvs within CAR T cells extends its potential applications across various therapeutic and biotechnological fields. In addition, the developed scFv library stands poised as a versatile resource, offering a rich foundation to unveil additional targets across diverse cancer types.

## Materials and Methods

### Isolation of PBMCs

Peripheral blood samples from 30 healthy donors were sourced from the German Red Cross Blood Bank, Heidelberg. Peripheral blood samples from 21 AML patients who had undergone donor lymphocyte infusion (DLI) were procured at Heidelberg University Hospital. Peripheral blood mononuclear cells (PBMCs) were isolated using Ficoll density gradient centrifugation in SepMate-50 tubes as described by the manufacturer. RNA extraction was performed with the RNeasy Mini Kit (Qiagen, Netherlands) according to the manufacturer’s instructions.

### Synthesis of cDNA

In brief, 500 ng isolated RNA template was used to synthesize the first strand cDNA with random hexamer primers and superScript IV reverse transcriptase (Thermofischer, USA) following the user manual.

### PCR amplification of antibody variable regions

A set of primers (supplemental Table S1) were used for amplifying the variable regions of antibodies. Individual forward and reverse primers were used for each gene. For each PCR reaction, 2 tubes of 50 µl reaction mixtures were prepared, containing 2.5 µl of cDNA. The amplification program consisted of an initial denaturation step at 98°C for 3 min, followed by 25-28 cycles of 98°C for 30 sec, annealing at 48-63°C for 30 sec, and extension at 72°C for 30 sec. The final extension was performed at 72°C for 3 min.

### Construction of scFv phagemid library

Amplified VH PCR products, 2 µg each, were individually subjected to double enzymatic digestion using NcoI (NEB, USA) and HindIII (NEB, USA), while VL PCR products were digested with MluI (NEB, USA) and NotI (NEB, USA) according to the manufacture’s user manual. The digested PCR samples were then purified from agarose gel using the Gel DNA recovery kit (Zymo Research, Germany). Purified VH PCR products were ligated with linearized pSEX81 (Progen, Heidelberg), digested with NcoI and HindIII, at a 4:1 mole ratio in the presence of T4 DNA ligase (Thermo Scientific, USA). Ligation products were concentrated by isopropanol precipitation and separately transformed into 25 µl electrocompetent *Escherichia coli* (*E.coli*)TG1 cells (BioCat, Germany) using a MicroPulser electroporator (Bio-Rad, CA) with the Ec1 program (1.8 kV, 1 pulse). Library size was quantified by performing serial dilutions of transformed cells and counting the number of clones. Subsequently, phagemid were extracted using the GeneJET plasmid Maxiprep kit (Thermo Scientific, USA). The VH-containing pSEX81 vector was digested with MluI and NotI and gel-purified for ligation with MluI/NotI double-digested VL PCR products.

### PacBio sequencing and bioinformatics analysis

scFv fragments were cut from constructed phagemids with NcoI and NotI and purified by agarose electrophoresis. The quantity and quality of purified scFv were measured by Qubit (Thermo Scientific, USA) and Bioanalyzer 2100 (Agilent, USA). scFv DNA fragments were then sequenced on the PacBio sequel II platform (Novogene, China). The sequencing coverage and quality statistics are summarized in Supplemental Table S2-S4. The high-fidelity reads obtained from PacBio sequencing were converted into a FASTA file and submitted to HighV-QUEST from IMGT^33^. The analysis was performed by choosing “homo sapiens”, “IG” receptor type and “Analysis of single chain Fragment variable”. “In-frame V(D)J” sequences were selected and further filtered with >85% identity between the V-REGION of the sequencing data and the V-REGION of the closest germline genes and alleles from the IMGT database.

### scFv-phage library production

25 ml overnight culture of *E.coli* transformants with scFv library were cultivated in 2xYT medium supplemented with 100 µg/ml ampicillin and 100 mM glucose. The culture was maintained at 37°C with shaking at 250 rpm till the optical density at 600 nm (OD_600_) of 0.5 (counting for 1.25 x 10^10^ cells) was reached. Then, 2.5 x 10^11^ plaque-forming units (pfu) of M13K07ΔpIII hyperphage (MOI=20) were introduced to the bacterial culture. The mixture was then incubated at 37°C without shaking for 30 min, followed by shaking at 200 rpm for an additional 30 min. Cells were pelleted by centrifugation and resuspended in 400 ml 2xYT medium supplemented with 100 µg/ml ampicillin and 50 µg/ml kanamycin. The suspension was cultured overnight at 25°C with shaking at 250 rpm. The culture supernatant was harvested and 100 ml of ice-cold 50% (w/v) PEG 6000 and 10 ml 4 M NaCl were added. The mixture was incubated on ice with gentle shaking for 2h. The phage particles were collected by centrifugation at 10,000 g for 2h at 4 °C and resuspended into 1 ml PBS containing 10% glycerol. The infective titers of the phage-scFv were determined using a plaque count assay. 10 ml *E. coli* XL1-Blue MRF’ cells were cultured in 2xYT medium at 37 °C up to OD_600_ of 0.5 and were exposed to a serial dilution of the phage for 30 minutes at 37°C. 100 µl of the samples were plated onto 2xYT agar plates containing 50 µg/ml kanamycin and incubated overnight at 37°C. The formed plaques were counted for calculating the phage titers.

### Panning of phage display scFv library

For the first panning round, 3 µg CLL1 antigen (AcroBiosystems, USA) per well in 100µl of PBS were immobilized on the Immuno 96 MicroWell Plates (Nunc, Denmark) overnight at 4°C. In the second and third panning rounds, 1µg and 0.3 µg of CLL1 were used respectively. After immobilization, antigen-containing wells were blocked with 5% (w/v) skim milk powder in PBS with 0.05% (v/v) Tween 20 (PBST). A phage library containing approximately 6 x 10^11^ phage particles in 50 µl panning block was then added to each well, followed by a 2-hour incubation at room temperature. Non-bound phages were removed by washing the plates 15 times with PBST followed by an additional 15 washes with PBS. The bound phages were eluted by incubating with 150 µl of 10 µg/ml trypsin for 30 minutes at 37°C. 150 µl exponentially growing *E. coli* TG1 cells (OD_600_ of 0.4-0.5) were added to the eluted phages with a 30-minute incubation at 37°C without shaking and 30 min at 37°C with shaking at 650 rpm in a block heater (Thermo Scientific, USA). Overall, 20 µl of the infected cells were then plated on 2xYT agar plates supplemented with ampicillin (100 mg/mL) and glucose (1% w/v) and incubated overnight at 37°C. For generation of selected scFv-phage for next round of panning, 720 µl 2xYT medium containing 100 µg/ml ampicillin and 100mM glucose was added to the rest 280 µl infected cells and incubated at 37°C with shaking to OD_600_ of 0.4-0.5. Bacterial cells were infected with 1 x10^10^ M13K07 helperphage particles (Thermo Scientific, USA) at 37 °C without shaking, followed by 30 min at 37 °C at 650 rpm. The MTP plate was centrifuged at 3220 g for 20 min. Medium was replaced with 1 ml 2xYT containing ampicillin and kanamycin, and incubated overnight at 30 °C and 850 rpm to produce new antibody phage. Phage-containing supernatant can directly be used for the next panning round. Single clones on the agar plates were picked up and phagemid DNA was sanger sequenced.

### Cloning of anti-CLL1 CAR

The coding sequences of the anti-CLL1 scFvs identified from panning were amplified via PCR. The purified PCR products were individually inserted into the Ncol/BamhI double digested retroviral vector RV-SFG.CD28.4-1BB.CD3 ζ (kindly provided by Prof. Malcolm Brenner, Center for Cell and Gene Therapy, Baylor College of Medicine, Houston, TX, USA) by using In-Fusion cloning (Takara Bio, Shiga, Japan) according to the manufacturer instructions. 1 µl of the recombinants was transformed into NEBstable competent cells (NEB, USA). Bacterial cells were spread on agarose plates, single clones were picked and the insertion of scFv was confirmed by Sanger sequencing.

### Generation of anti-CLL1 CAR-T cells

Anti-CLL1.CD28.4-1BB.CD3ζ retroviruses were produced in HEK293T cells with co-transfected with 3.75 µg packaging plasmid PegPam3, 2.5 µg envelope plasmid RDF (both provided by Professor Malcolm Brenner), and 3.75 µg transfer plasmid RV-SFG.CLL1.CD28.4-1BB.CD3ζ. Cryopreserved PBMCs were thawed, and 1×10^6^ cells were seeded onto each well of 24-well plates coated with anti-CD3/anti-CD28 antibodies. 24 hours later, cells were stimulated with interleukin-7 (IL-7) and interleukin-15 (IL-15). On the third day, 0.2 x10^6^ activated T cells were added to each well of 24-well plates coated with 1 ml retrovirus CLL1.CD28.4-1BB.CD3ζ collected at 48h and 72h. The efficiency of transduction was assessed on the fourth day following transduction via flow cytometry using an anti-human goat F(ab) IgG (H+L) PE antibody (Dianova, Hamburg, Germany). The quantity of effector cells employed for functional assays was adjusted based on the determined transduction efficiency. For the cultivation of CAR T cells, a culture medium consisting of 45% RPMI-1640 and 45% EHAA media (Clicks) supplemented with 2 mM L-glutamine, 10% fetal bovine serum (FBS), 10 ng/ml IL-7, and 5 ng/ml IL-15 were used.

### Cell lines

Cell lines MV4-11 (RRID: CVCL_0064), HL60 (RRID: CVCL_0002) and HEK 293T (RRID: CVCL_0063) were obtained from American Type Culture Collection. K562 (RRID: CVCL_0004) cells were purchased from the German Collection of Microorganisms and Cell Cultures (Braunschweig, Germany). ZsGreen-MV4-11 was obtained by lentiviral transduction of ZsGreen gene into wide type MV4-11 cells. All leukemic cell lines were cultured in RPMI-1640 media supplied with 10% fetal bovine serum (10% FBS) and 1% of penicillin and streptavidin in a humid incubator with 5% CO_2_ at 37°C. HEK 293T cells were cultured in IMDM medium containing 10% FBS. All cell lines have been authenticated using SNP profiling within the last three years and are mycoplasma-free.

### Surface protein analysis

The quantification of CLL1 and CD33 expression on tumor cell surface, and CD3 expression on T cells was conducted through flow cytometry using anti-CLL1-APC (clone 50C1), anti-CD33-APC (clone p67.7) and anti-CD3-BV510 (clone OKT3), respectively. All antibodies were purchased from Biolegend, Netherland. To ensure the accuracy of these staining procedures, dead cells were effectively excluded using the 7AAD dye (BD Biosciences, USA).

### Intracellular cytokine staining

To evaluate cytokine release, CAR T cells were cultured alongside target tumor cells HL60, MV4-11, MV4-11 zsGreen at a ratio of 2:1 for 5 hours in 96-well plates (Sarstedt, Germany) in the presence of 5 µg/ml Brefeldin A (Biolegend, CatNo: 42060, Netherland) and 2.0 µM Monensin (Biolegend, CatNo: 420701, Netherland) for 5 hours. Cells were then fixed and permeabilized using the FoxP3 buffer set (Miltenyi Biotec, Bergisch Gladbach, Germany). Intracellular interferon γ (IFNγ) and tumor necrosis factor α (TNFα) were stained with anti-IFNγ-Alexa Fluor 488 (clone 4S.B3, Biolegend, Netherland) and anti-TNFα-BV421 antibodies (clone Mab11, Biolegend, San Diego, USA). All experimental samples were analyed using LSRII or Symphony A3 cell analyzer (BD Biosciences, USA) and the resulting data were processed and interpreted using FlowJo software.

### Long-term coculture assay

2 x 10^4^ anti-CLL1 CAR T cells were cocultured with 4 x 10^4^ HL60, MV4-11 and MV4-11 zsGreen, cells in a 48-well flat-bottom plate containing 800 µl of CAR T cell culture medium (see above). Every second day, additional 4 x 10^4^ tumor cells were introduced into the coculture to rechallenge the CAR-T cells. CAR T cells and tumor cells were monitored via flow cytometry.

### In mice experiments

Six to ten-week-old NOD.Cg-Prkdcscid IL2rgtm1Wjl/SzJ (NSG) mice were obtained from the German cancer research center (DKFZ) in Heidelberg and housed at Interfaculty Biomedical Facility (IBF). Following a 10-day acclimatization period, each mouse received an injection of 1× 10^6^ tumor cells via the tail vein for establishment of AML xenografts. In the experimental groups, 5 days post-initiation, each mouse received 5× 10^6^ CAR T cells via a tail vein injection. Tumor burden and trafficking were assessed through bioluminescent imaging (BLI) using the Xenogen In Vivo imaging System (IVIS; Caliper Life Sciences, USA). All experimental procedures were conducted in accordance with the guidelines outlined by IBF, Heidelberg.

### Statistical Analysis

Statistical analyses were conducted using GraphPad Prism 9 software (GraphPad Software, La Jolla, CA, USA). The calculation of p-values was carried out with the unpaired parametric t-test.

## Results

### scFv phagemid library generation

The VH and VL genes were PCR-amplified from 30 healthy donors and 21 AML patients in complete remission after donor lymphocyte infusion (Figure 1A). Initially, recombination-based In-fusion cloning was used to insert these genes into the phagemid vector pSEX81, but overlapping PCR failed to generate VH-(G4S)3-VL fragments when multiple VH and VL fragments were introduced into the reaction simultaneously (Supplemental Figure S1A). A switch to the traditional cloning with restriction enzyme digestion and ligation was made (Supplemental Figure S1B). Individual unbiased PCR reactions, each with a unique forward primer and a corresponding reverse primer, yielded expected ∼400 bp bands on agarose gel (Supplemental Figure S2 A, B). 24 VH fragments digested with NcoI/HindIII ligated into pSEX81 vector and transformed into TG1 bacterial cells. Approximately 0.51 x 10^6^ colonies were obtained from each transformation, resulting in a total of ∼12 x 10^6^ colonies. Then, the pSEX81 vector carrying the VH repertoire was digested with MluI/NotI for insertion of VL genes. A total of 264 transformations were performed, yielding ∼1.35 x 10^8^ colonies (Figure 1A, Supplemental Figure S1).

**Figure 1.**
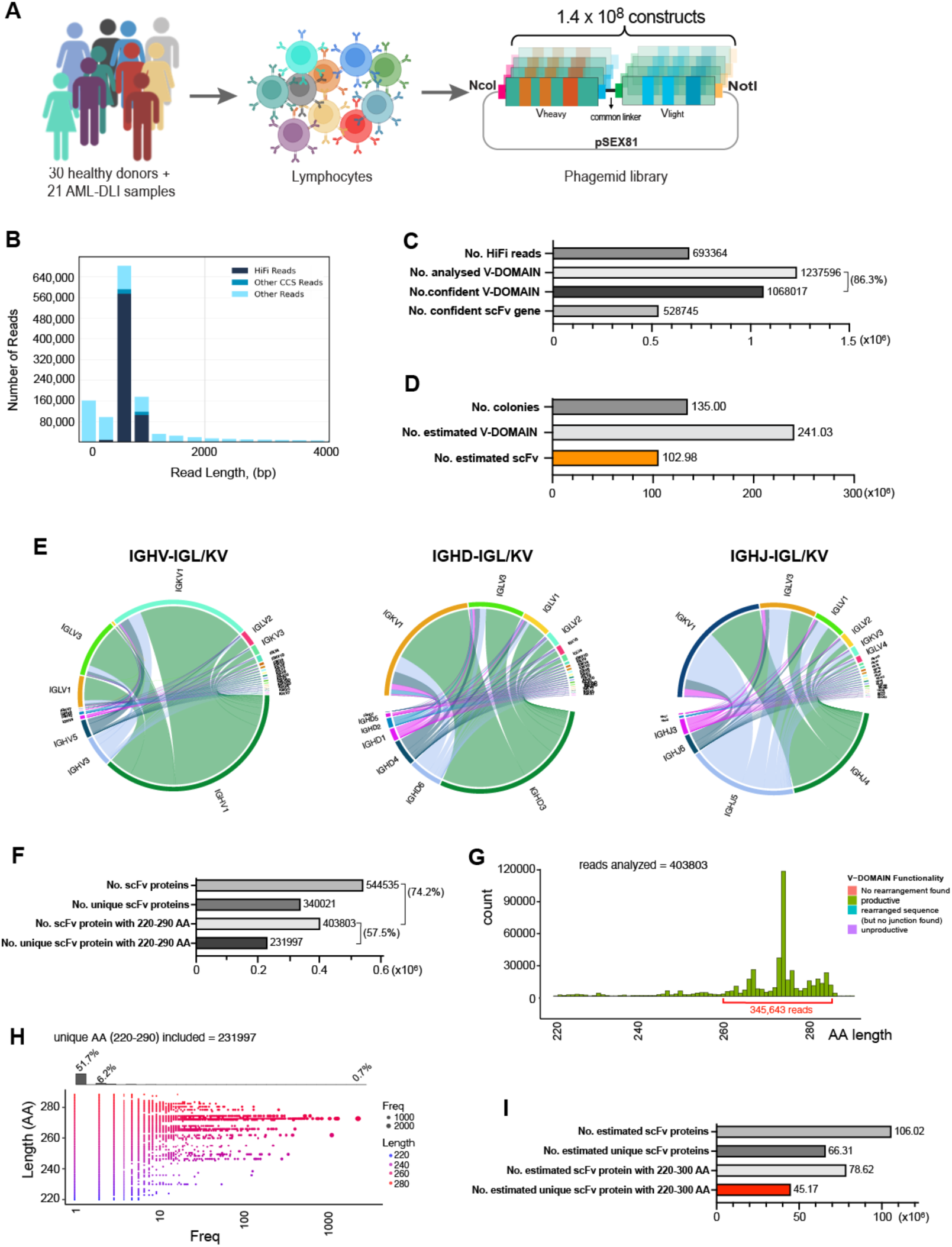
High-throughput sequencing and data analysis of the scFv phagemid library. (A) Brief diagram depicting the scFv phagemid library generation. (B) PacBio read length distribution using one SMRT cell. HiFi reads (QV≤20) in dark blue, other CCS Reads (three or more passes, but QV < 20) in blue, and other reads in light blue. (C) Identified variable domains and scFvs with the PacBio HiFi reads. (D) Estimation of variable domains and scFvs in the generated phagemid library. (E) Chord diagram illustrating the associations and frequency of variable (left), diversity (middle) and joining (right) genes of the heavy chains with the variable genes of the light chains. (F) Identified scFv proteins with the PacBio HiFi reads. (G) Amino Acids (AA) length distribution. (H) Read frequency of unique scFv with 260-290 AA long. (I) Calculation of scFvs proteins in the generated phagemid library

### High-throughput sequencing and analysis of scFv phagemid library

To ensure scFvs for sequencing truly representing the phagemid library, scFv fragments were directly excised from phagemids, yielding ∼800 bp DNA (Supplemental Figure S3A, B). From one PacBio Sequel II SMRT cell, ∼2.7 x 10^8^ subreads were generated, resulting in ∼1,5 x 10^6^ circular consensus sequence (CCS) reads (Supplemental Figure S3 C and Table S2-3). CCS reads were further filtered to eliminate instances of double-loaded wells or any reads with low accuracy. Following filtration, 693,364 long reads with high accuracy, termed HiFi reads, were obtained (Figure 1B, Supplemental Table S4). The quality value (QV) of these HiFi reads is equal to or greater than 20 (QV ≥ 20), representing an accuracy of 99% or higher. The average length of the HiFi reads was 805 bp.

Mapping these HiFi reads to human germline IgG gene segments with HighV-QUEST from IMGT identified ∼1.2 x 10^6^ variable domains (i.e., one scFv contains two V-REGIONs) were identified (Figure 1C). Of note, 1,068,017 (86.3%) V-REGIONs showed >85% identity, which is the standard filter threshold of antibody repertoire, to the V gene of closest antibody in the IMGT database and were defined as “confident” sequences. Among them, 99% represented scFv clones (528,745) containing both V domains of heavy and light chains (Figure 1C).

Through re-evaluating the phagemid library which contains ∼1.35 x 10^8^ colonies, based on the PacBio sequencing statistics, approximately 1.03 x10^8^ scFvs were identified, covering nearly all human germline IgG subfamilies (Figure 1D). Detailed analysis revealed, without considering alleles, 56 variable genes (HV), 25 diversity genes (HD), and 6 joining genes (HJ) for heavy chains; 26 variable genes (KV) and 5 joining genes (HJ) for *Kappa* light chains; and 38 variable genes (LV) and 6 joining genes (LJ) for *lambda* light chains (Supplemental Figure S4). All identified gene segments are functional. The associations and frequencies of variable, diversity and joining genes in the heavy chains with variable genes in the light chains vary. Predominant gene associations include IGHV1 and IGKV1 (41.3%), IGHV1 and IGLV3 (17.4%); IGHD3 and IGKV1 (34.7%), IGHD3 and IGLV3 (14.7%); IGHJ4 and IGKV1 (22.0%), IGHJ5 and IGKV1 (21.2%), IGHJ4 and IGKV3 (9.3%), IGHJ5 and IGKV3 (9.2%) (Figure 1E).

### Amino acid distribution of scFvs

Biostrings was used to translate DNA information into amino acids (AA). A total of 544,535 translation products were found and 403,803 (74.2%) have length of 220-290 AA indicating full-length scFv proteins, and 85.6% (345,643) fell within the 260-285 AA range (Figure 1F, G). The most dominant length was 274 AA, accounting for 2.3% of the total (Figure 1G). Among the 403,803 full-length scFvs, 231,997 (57.5%) were unique. The most frequently observed scFv appeared 2,881 times (0.7%), while the lowest read scFvs, totaling 208,961, made up 51.7% of the entire library (Figure 1H). Recalculation of AA diversity in the phagemid library, estimated at ∼1.35 x 10^8^ colonies, indicated ∼1.0 x10^8^ scFv proteins with length of 220-300 AA, with ∼4.5 x 10^7^ being unique (Figure 1I).

### Discovery of anti-CLL1 scFvs

As a proof of principle, we identified scFvs targeting CLL1. The scFv-expressing phage library was packaged in TG1 bacterial cells, confirmed by the 95 kD scFv-pIII fusion protein on the western blot gel (Figure 2A). The scFv library was then screened against purified CLL1. CLL1 antigen was chosen due to its high expression on >90% malignant cells compared to healthy stem cells ^34^ (Figure 2B). We performed three independent panning experiments, each consisting of three rounds of selection. After the 3rd selection of each panning experiments, 35, 25 and 36 colonies were randomly selected for sanger sequencing, resulting in the identification of 11, 18, 12 functional full-length scFvs, with 3, 3, and 5 being unique in each panning, respectively (Figure 2C). In total, six unique scFv sequences were identified and used for further analysis (Figure 2C).

**Figure 2.**
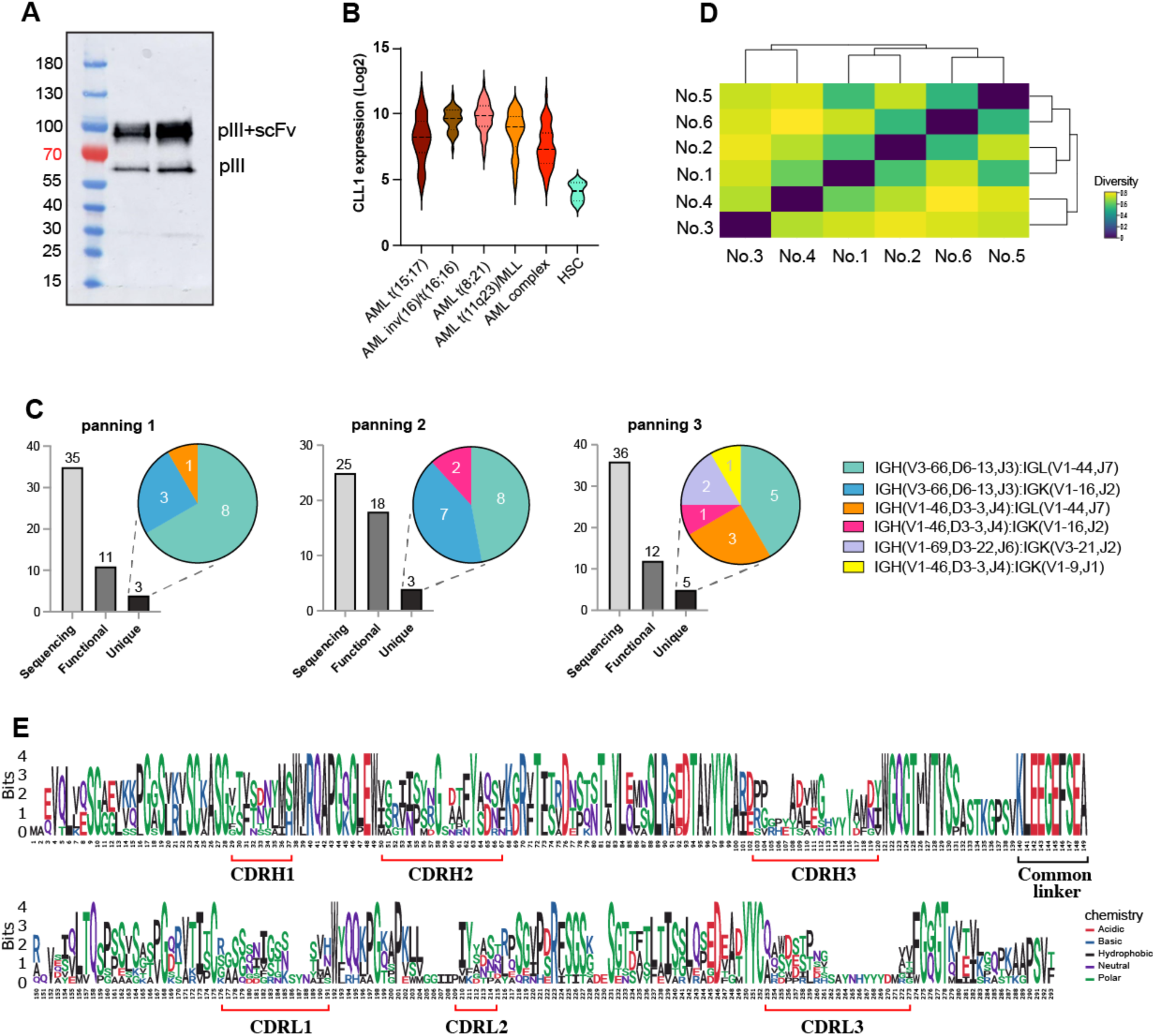
Identification of scFvs with specific affinity to antigen CLL1. (A) Western Blot of packaged phage. Upper band indicated pIII-scFv fusion protein. (B) CLL1 expression in different AML and healthy stem cells. Data presented are from Bloodspot. (C) Unique scFvs identified from 3 independent panning experiments, each consisting of 3 rounds of selections. (D) Heat map represents the amino acid conservation among the identified scFv sequences. The function calculates an identity matrix of pairwise distances from aligned sequences, containing the square root of these distances. For example, the sequence identity between No.4 and No.6 is 43.1%. Then, the square root of (1.0-0.431), 0.7543, is represented according to the color bar. (E) Sequence alignment of identified unique scFvs.

Amino acid conservation analysis revealed sequence identities ranging from 43.1% (between No.4 and No.6) to 75.5% (between No.5 and No.6) (Figure 2D). To explore sequence patterns, sequence logos of these six unique scFv sequences were generated with ggseqlogo. As expected, the complementarity-determining regions (CDR) showed the highest variability, while the framework regions (FR) scaffolds were more conserved (Figure 2E). Moreover, the linker peptide KLEEGEFSEA between the heavy and light chains remained constant, further supporting the reliability of sequences (Figure 2E).

### CLL1-CAR T cell generation and characterization

Each identified scFv was individually inserted into the third generation CAR vector containing expression cassettes for a human IgG1 CH2CH3-derived hinge, the stimulatory domains of CD28 and 4-1BB, and the CD3ζ signaling domain (Figure 3A). Anti-CLL1-CAR T cells were generated by retroviral transduction. CAR expression levels in T cells transduced with scFv number 1, 2 and 4 reached 24.2% (range: 20.5% - 37.9%), 36.8% (range: 30.7% - 41.2%) and 12.6% (range: 8.9% - 15.5%), respectively, while construct 3 showed no CAR expression. T cells carrying constructs 5 and 6 displayed higher CAR expression levels of 48.9% (range: 39.8% - 51.6%) and 65.1% (range: 54.3% - 68.3%). (Figure 3B). In subsequent assays, T cells numbers were adjusted based on the percentage of CAR-positive cells.

**Figure 3.**
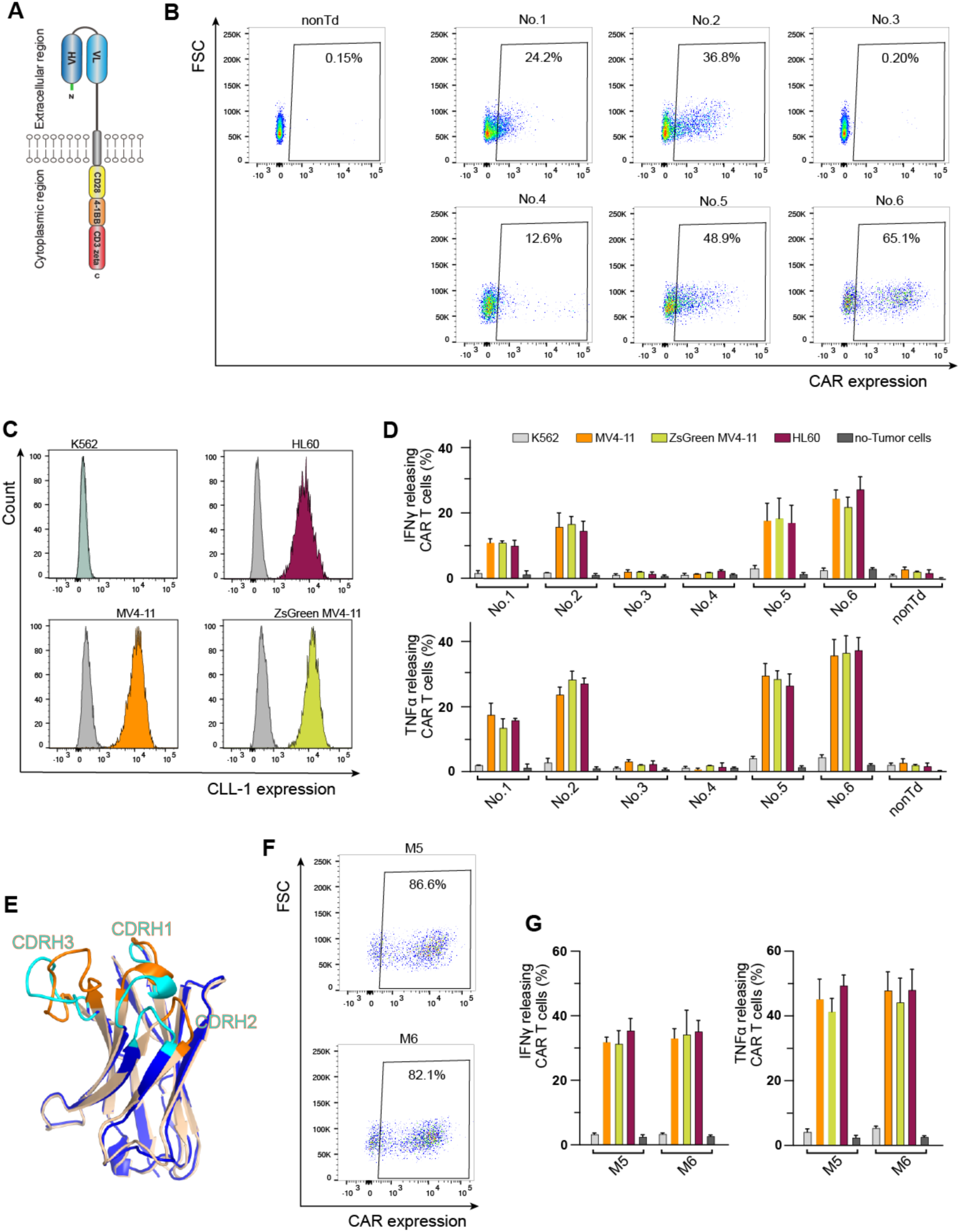
CAR T cells carrying scFvs identified from panning. (A) Schematic diagram of the third generation CLL1-targeting CAR construct. (B) Representative CAR expression of T cells individually transduced with CAR constructs carrying scFv number 1-6 identified from panning. nonTd: nontransduced T cells. T cell donors, n = 3. (C) CLL1 expression in different cell lines. Unstained cells are shown in grey. (D) Intracellular cytokine secretion of T cells individually transduced with CAR constructs carrying scFv number 1-6 identified from panning, upon stimulation of tumor cells or without stimulation (dark grey). nonTd: nontransduced T cells. Experiments were conducted in triplicates. Mean values were calculated for each group; error bars indicate SD. (E) 3D structure alignment of heavy chains of scFvs identified through panning with our generated scFv library (wheat) and scFv h6E7.L4H1e (WO2016/205200) (blue). Wheat: heavy chain of identified No.5 and 6, sharing identical sequence, with CDRs highlighted in cyan. Blue: reported heavy chain, with CDRs highlighted in Orange. Structures were predicted using Alphafold google colab. (F) Representative CAR expression of T cells carrying modified scFvs. T cell donors, n = 3. (G) Intracellular cytokine secretion of T cells individually transduced with modified CAR constructs. Experiments were conducted in triplicates. Mean values were calculated for each group; error bars indicate SD.

To assess the functionality and specificity of the generated CAR T cells, CLL1 expression was first evaluated across various leukemia cell lines. K562 cells showed no CLL1expression, whereas MV4-11, ZsGreen MV4-11, and HL60 cell lines were positive for CLL1 (Figure 3C). Intracellular cytokine IFNγ and TNFα secretion of CAR T cells expressing scFvs 1, 2, 5 and 6 was detected upon stimulation with CLL1-expressing MV4-11, ZsGreen MV4-11, and HL60 cell lines (Figure 3D, Supplemental Figure S5). Notably, no cytokine secretion was detected in CAR T cells co-cultured with CLL1-negative K562 cells. In addition, T cells transduced with non-CAR constructs (3 and 4) and non-transduced T cells (nonTd) did not release cytokines under the same condition (Figure 3D).

Attempts to enhance CAR expression of identified constructs to levels comparable to our previous anti-CD33 CAR (> 80%) yielded limited improvement. We then compared our identified anti-CLL1 scFvs with existing counterparts. We discovered that the heavy chain sequence of constructs 5 and 6 (which are identical) showed 81.97% identity to the reported sequence h6E7.L4H1e (Patent number: WO2016/205200), particularly in the FR regions (∼93% identity), while the CDRs varied widely (Figure 3E). Then, we incorporated the published heavy chain CDR sequences into our constructs while retaining the original light. These modified CAR constructs showed a marked increase in transduction efficiency, with rates reaching 86.6% (range: 75.9% - 89.2%) for M5 and 82.1% (range: 71.9% - 85.6%) for M6 (Figure 3F). Moreover, IFNγ and TNFα secretion was also significantly increased (Figure 3G).

### *In vitro* Cytotoxicity of CLL1-CAR T cell

The most effective CAR T cells, characterized by over 50% CAR-positive cells and high levels of cytokine secretion (i.e. constructs No.6, M5 and M6), were selected to assess their tumor-killing capacity. First, a short-term 2-day coculture experiment showed that these CAR T cells effectively eradicated CLL1-positive tumor cells MV4-11, but having no impact on CLL1-negative K562 cells (Figure 4A). Notably, K562 cells, which are CD33-positive, could be targeted and destroyed by CD33-directed CAR T cells and stimulated these CAR T cell to release cytokines (Supplemental Figure S6). These observations indicate the specificity of these anti-CLL1 CAR T cells, which selectively target tumor cells expressing the CLL1 antigen.

**Figure 4.**
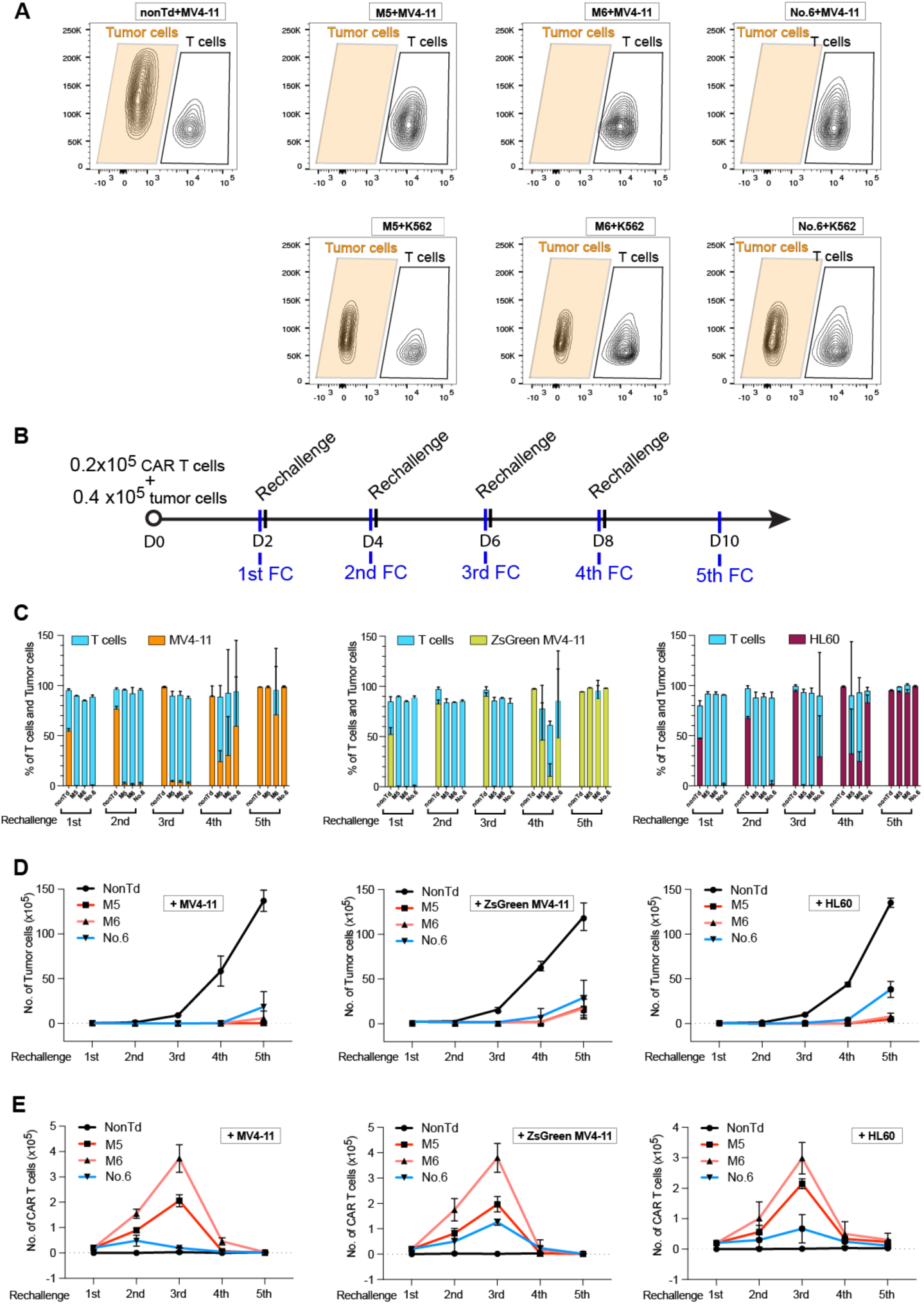
Cytotoxicity of selected CAR T cells, carrying scFv No.5 identified through panning with our generated scFv library, and modified scFv M5 and M6, with reported CDRs. (A) two days of short-time killing assay. Tumor cells (T) and CAR T cells (E) were cocultured at an E:T ratio of 1:2 for 2 days and then analyzed by FC. The CLL1-positive MV4-11 tumor cells and CLL1-negative K562 tumor cells were used in this assay. T cell donors, n = 3. Data is show from donor 1. (B) Work-flow of long-term coculture assay. Eight wells were prepared for each coculture, each containing CAR T cells (T cell donor, n=3) and one type of tumor cells. Every second day, cells from one well were collected for FC analysis. CAR T cells in the remaining wells were rechallenged with fresh tumor cells. (C) Percentage of T cells and tumor cells in the long-term coculture at the time point of FC analysis. Experiments were conducted in triplicates using CAR T cells generated from three healthy donors. Mean values were calculated for each group; error bars indicate SD. The high heterogeneity among T cell donors resulted in significant variation in SD for the last rounds of killing. (D) Tumor cell expansion during the long-term coculture. (E) CAR T cell expansion during the long-term coculture.

Next, the long-term cytotoxicity of these CAR T cells was evaluated using a prolonged coculture assay (Figure 4B). During the initial phase, CAR T cells successfully eliminated CLL1-positive tumor cells (MV4-11, ZsGreen MV4-11 and HL60). Additional tumor cells were added into the culture every second day to rechallenge the CAR T cells. In this analysis, all CAR T cells displayed complete elimination of tumor cells for three rounds (in the condition without addition of cytokines IL7/15). However, during the fourth rechallenge, a proportion of tumor cells persisted. By the fifth round, their numbers had increased to over 95% (Figure 4C).

The expansion of both tumor cells and CAR T cells was tracked throughout the long-term coculture assay. Tumor cells in the non-transduced T cell groups expanded rapidly, increasing from the initial 0.04 x10^6^ to over 10 x10^6^. In contrast, tumor cell growth in the CAR T cell-treated groups was markedly suppressed, with noticeable inhibition occurring after the fourth rechallenge (Figure 4D). The proliferation of CAR T cells peaked during the third rechallenge, expanding approximately 20-fold, from the initial 0.02 x10^6^ to about 0.4 x10^6^, demonstrating robust expansion of CAR T cells(Figure 4E).

### *In vivo* Cytotoxicity of CLL1-CAR T cell

We further conducted an *in vivo* cytotoxicity experiment using NSG mice. Initially, 1×10^6^ ZsGreen-expressing tumor cells were injected into mice. Five days later, after confirming tumor cell engraftment, 5×10^6^ CAR T cells (constructs No.6, M5 and M6) or non-CAR transduced T cells were injected into the mice. Three days post T cell injection, mice received CAR T cells No.6 and M5 developed severe illness and were euthanized. In contrast, mice treated with CAR T cells M6 or non-transduced T cells remained normal. Tumor cells progression in these groups were monitored weekly using BLI (Figure 5A). At 7 days post-CAR T cell injection, a reduction in tumor cells was observed in the CAR T cell-treated mice (Figure 5B, C). By day 14, three mice from the non-transduced T cell group died, while all six mice in the CAR T cell-treated group survived, displaying markedly lower tumor cell signals (Figure 5C).

**Figure 5.**
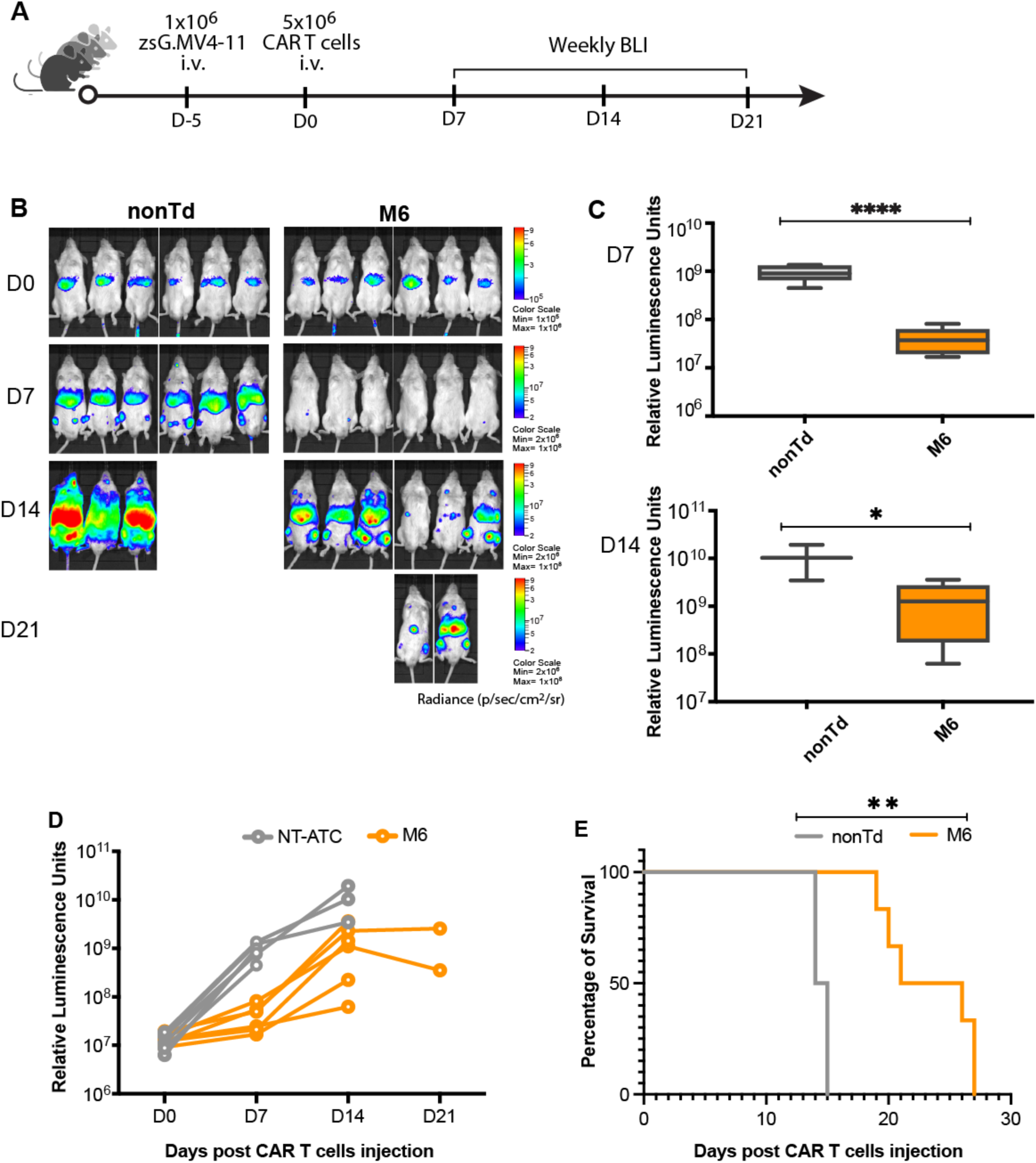
in vivo Cytotoxicity. (A) Work-flow of in vivo killing assay. NSG mice engrafted with zsGreen MV4-11 were treated with nontransduced T cells (nonTd) (n=6) and anti-CLL1 CART cells (n=6). Tumor regression was measured by bioluminescent imaging (BLI) weekly. (B) Tumor burden in mice on the day of CAR T cell injection, 7, 14, and 21 days post CAR T injection as measured by BLI. (C) Tumor cell luminescence signal 7 days post CAR T injection (upper) and CAR T and 14 days post CAR T injection (lower). *: p < 0.05; ****: p < 0.0001. (D) Overall tumor cell luminescence signal along experimental period. (E) Survival possibility of mice in nontransduced T cells (nonTd) and anti-CLL1 CART cells.

Pertaining to the overall tumor regression observed in mice, the CAR T cell-treated group exhibited lower tumor burden compared to the non-transduced T cell group (Figure 5D). Survival in the CAR T cell-treated group was prolonged compared to the non-transduced T cell group. In the control group, mice died on days 14 and 15. Mice in the CAR T cell-treated group started to succumb to disease on day 19, with the longest survival day extending to day 27 (Figure 5E).

## Discussion

Personalized immunotherapy requires rapid and patient-specific targeting of tumor cells. Antibodies and scFvs exhibit complexity through their composition of heavy and light chains, housing variable regions crucial for antigen recognition. These highly diverse variable regions are generated through random recombination of V(D)J gene segments. According to the V Base^35^ and IMGT databases, there are 56 VH, 26 DH, 6 JH, 42 VK, 5 JK, 38 VL and 6 JL genes, reflecting the inherent complexity and diversity. Constructing scFv libraries is significantly challenging. In this study, early efforts employed recombination-based In-Fusion cloning to generate the scFv phagemid library, which avoid enzymatic digestion of the variable regions. However, the main challenge was the efficient and simultaneous amplification of multiple DNA fragments via overlap PCR to assemble VH-(G4S)3-VL fragments. Consequently, we adopted a final approach using restriction enzymatic digestion and fragment ligation. To reduce potential PCR bias and enhancing the library’s diversity, we designed specific primers with minimized wobble nucleotides, covering the entire breadth of functional antibody gene repertoires. Moreover, each forward primer was independently used in PCR reactions to amplify their specific genes.

Quality assessment of the scFv library is crucial for ensuring the identification of optimal scFvs that are specific to chosen antigens. Sequencing technology plyas a pivotal role in determining the depth, accuracy and integrity of the analysis. Many NGS platforms face limitations with scFv sequences, which typically span 800 bp. Previous studies used NGS methods to capture reads spanning only one single variable domain or the CDRs resulted in the loss of crucial information embedded in scFv sequences, specially, the association of the two distinct VH and VL domains ^36–38^. Thanks to the technological development, the newly emerging third-generation PacBio sequencing is capable of generating long reads with high fidelity. We used PacBio Sequel II and yielded 6.9 x 10^5^ high-accuracy HiFi reads from one single SAMT cell. IMGT analysis revealed a rich diversity of 1.07 x 10^6^ confident scFv sequences. This diverse repertoire encompassed antibody gene segments from almost all functional subfamilies of variable heavy and light chains, exhibiting a broad representation of VH and VL gene diversity within the library. Translating these DNA sequences into amino acids unveiled a considerable number of full-length scFv proteins. When extrapolated to the scFv phagemid library, it was estimated to contain approximately 4.5 x 10^7^ unique full-length scFvs. Our current library was estimated to have high diversity and coverage. However, dominance of certain subgroups was observed. Based on these assessment outcomes, an optimization of the scFv library, for instance by using a greater diversity of donors, or adjusting the subfamily composition, can enhance the distributions of antibody subfamilies and increase the proportions of full-length scFvs.

The functional characterization of scFv library validated the potential to obtain therapeutically active scFvs. Rigorous screening in this study identified scFvs with potential binding to the CLL1 antigen. Integration of these scFv sequences into the CAR backbone vector, which was used in our previous CD33-directed CAR project and in the ongoing clinical *CD19-directed CAR T cell therapy* at the Heidelberg University Hospital ^7,39^ (NCT03676504), facilitated the generation of CLL1-specific CAR T cells. Subsequent analyses revealed variable levels of CAR expression across different constructs. Integration of reported CDR sequences (Patent NO. WO2016/205200) into our scFvs showed improved CAR expression. Further functional validation demonstrated the specificity and potency of CAR T cells carrying selected and modified scFvs, revealing specific killing and cytokine release stimulated by CLL1-positive tumor cells *in vitro*. However, in-depth investigations revealed challenges pertaining to sustaining long-term efficacy. *In vitro* long-term coculture assays observed persistent tumor cells after successive multiple rechallenges. These findings hinted at potential limitations in sustained CAR T cell efficacy against tumor cells over prolonged exposures. Further investigations are warranted to elucidate mechanisms leading to the loss of CAR T cell potency over time and to enhance their long-term persistence and efficacy. Nevertheless, *in vivo* experiments conducted in NSG mice further substantiated the efficacy of these CAR T cells, displaying diminished tumor cell signals and improved survival rates in mice treated with CAR T cells compared to controls. This highlighted the promising therapeutic potential of the generated CAR T cells against CLL1-positive tumors in animal models.

In summary, this study’s generation of a proprietary scFv phage display library using samples from both healthy donors and AML patients in complete remission, combined with thorough profiling, alongside the successful discovery and validation of effective CAR T cells, present valuable pathways for the development of immunotherapeutic approaches against cancer. With the established scFv library, and the upcoming improved version, further targets could be explored for identification of specific scFvs for therapeutic purposes.

## List of abbreviations in alphabetical order

2YT: 2x yeast extract tryptone medium
AA: amino acid
AML: acute myeloid leukemia;
BLI: bioluminescent imaging
CAR: chimeric antigen receptor;
CAR T cells: Chimeric antigen receptor T cells;
CCS: circular consensus sequence;
CD: cluster of differentiation;
cDNA: complementary deoxyribonucleic acid;
CDR: complementarity-determining region
CLL1: C-35 type lectin-like molecule-1;
*E.coli*: *Escherichia coli* bacterial cells;
FACS: fluorescence-activated cell sorting;
FBS: fetal bovine serum;
FC: flow cytometry;
FDA: U. S. Food and Drug Administration;
FLT-3: fms-like tyrosine kinase 3;
FLT3-L: fms-like tyrosine kinase 3 ligand;
FR: framework regions;
G-CSF: granulocyte colony stimulating factor;
HD: diversity genes from heavy chains;
HJ: joining genes from heavy chains;
HV: variable genes from heavy chains;
ICS: intracellular cytokine staining;
IG: immunoglobulin;
IFN: interferon;
IL: interleukin
IMDM: Iscove’s Modified Dulbecco’s Medium;
IMGT: the international ImMunoGeneTics information system;
KD: diversity genes from *Kappa* light chains;
KV: variable genes from *Kappa* light chains;
LD: diversity genes from *lambda* light chains;
LV: variable genes from *lambda* light chains;
OD_600_: optical density at 600 nm
PacBio: Pacific Biosciences technology
PBMC: peripheral blood mononuclear cell;
PBS: phosphate-buffered saline;
PBST: phosphate-buffered saline with Tween 20;
PCR: polymerase chain reaction;
RNA: ribonucleic acid
SCF: stem cell factor;
scFv: single chain variable fragment;
TNF: tumor necrosis factor;
TPO: thrombopoietin.
VH: the variable regions of the antibody’s heavy chain
VL: the variable regions of the antibody’s light chain

## Acknowledgements

We thank Prof. Patrick Most, Dr. Eric Meinhardt and Dr. Martin Busch for providing the Biorad Micropulse Electroporator. Special thanks to Michael Heß for collecting and providing the AML-DLI patient samples.

## Contributions

Y.L. and C.M.-T. conceived the study, designed the strategies and wrote the original manuscript. Y.L. wrote the protocols, conducted most of the experiments, guided A.L., J.B. and E.C., analyzed data and created the figures. D.S. performed the mice experiments and analyzed the data. A.L. and E.C. generated the scFv phagemid library. A.L. and J.B. performed panning. J.B. generated the ZsGreen MV4-11 cell line. C.L. analyzed selected scFv sequences. M.M. provided the script for chord diagram. T.L. provided patient samples. H.Y. analyzed PacBio sequencing data. M.S. introduced CAR T cell experiments. T.M. and F.Z. participated in the discussion of the project. F.Z. provided insights into figure generation. C.R. analyzed PacBio sequencing data. C.M.-T. supervised and directed the project. All authors have read and commented the manuscript.

## Data Availability Statement

All data supporting the finding of this study can be obtained from the corresponding authors upon reasonable request.

## Ethics Statement

Primary sample collection and analysis were approved by the Ethics Committee of the University of Heidelberg (S-686/2018). Informed written consent was obtained from all patients before experiments. All experiments were conducted in compliance with the Declaration of Helsinki.

