## supplemental information for "Integrated scFv identification and CAR T cell generation for AML targeting in vivo"

**This supplementary includes:**

Supplemental figure S1: Overview of cloning approach used in this study
Supplemental figure S2: PCR products of VH and VL genes
Supplemental figure S3. High-throughput sequencing of the scFv phagemids library
Supplemental figure S4: Frequency and distribution of subfamilies and genes of heavy chains and light chains from the PacBio sequencing
Supplemental figure S5: ICS assay gating approach
Supplemental figure S6: K562 cells are antiCD33.CAR T cells sensitive

Supplemental table S1. Primers used in this study for amplifying VH and VL genes
Supplemental table S2. Subreads statistics of PacBio sequencing
Supplemental table S3. CCS reads statistics of PacBio sequencing
Supplemental table S4. HiFi reads statistics of PacBio sequencing

**Supplemental Figure S1**

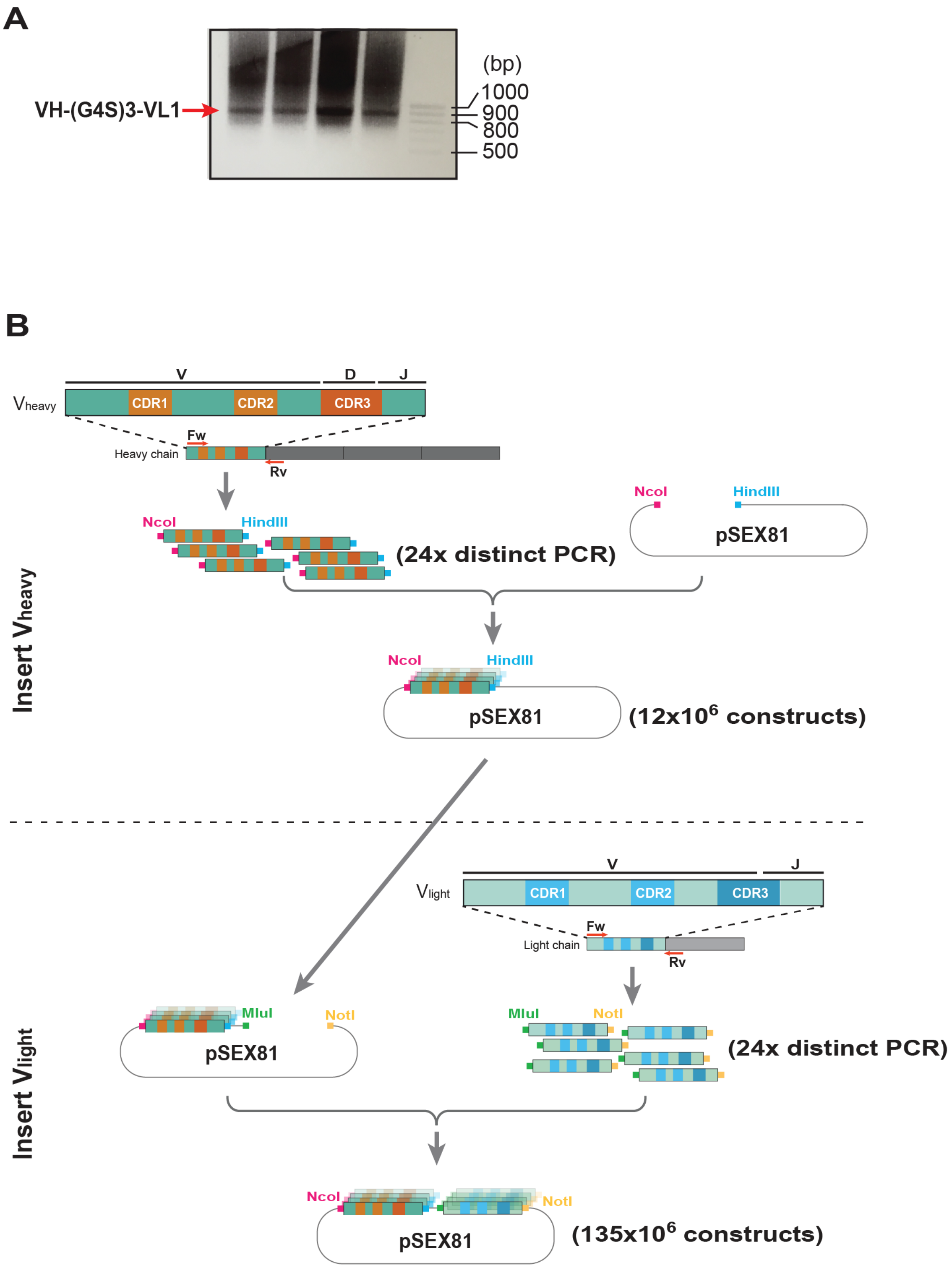

**Figure S1**. Overview of cloning approach used in this study. (A) Electrophoresis agarose gels of the overlap PCR attempting to assemble VH-(G4S)3-VL fragments. Successful amplification was achieved when single VH and single VL DNA fragment were used as the templates (Supplemental Figure S1A). However, when multiple VH and VL DNA fragments were introduced into the reaction simultaneously, the amplification of VH-(G4S)3-VL fragments proved to be unattainable. Desired bands were not consistently obtained. (B) Overview of digestion-ligation cloning approach used in this study, involving the sequential insertion of the heavy chain genes followed by the light chain genes.

**Supplemental Figure S2**

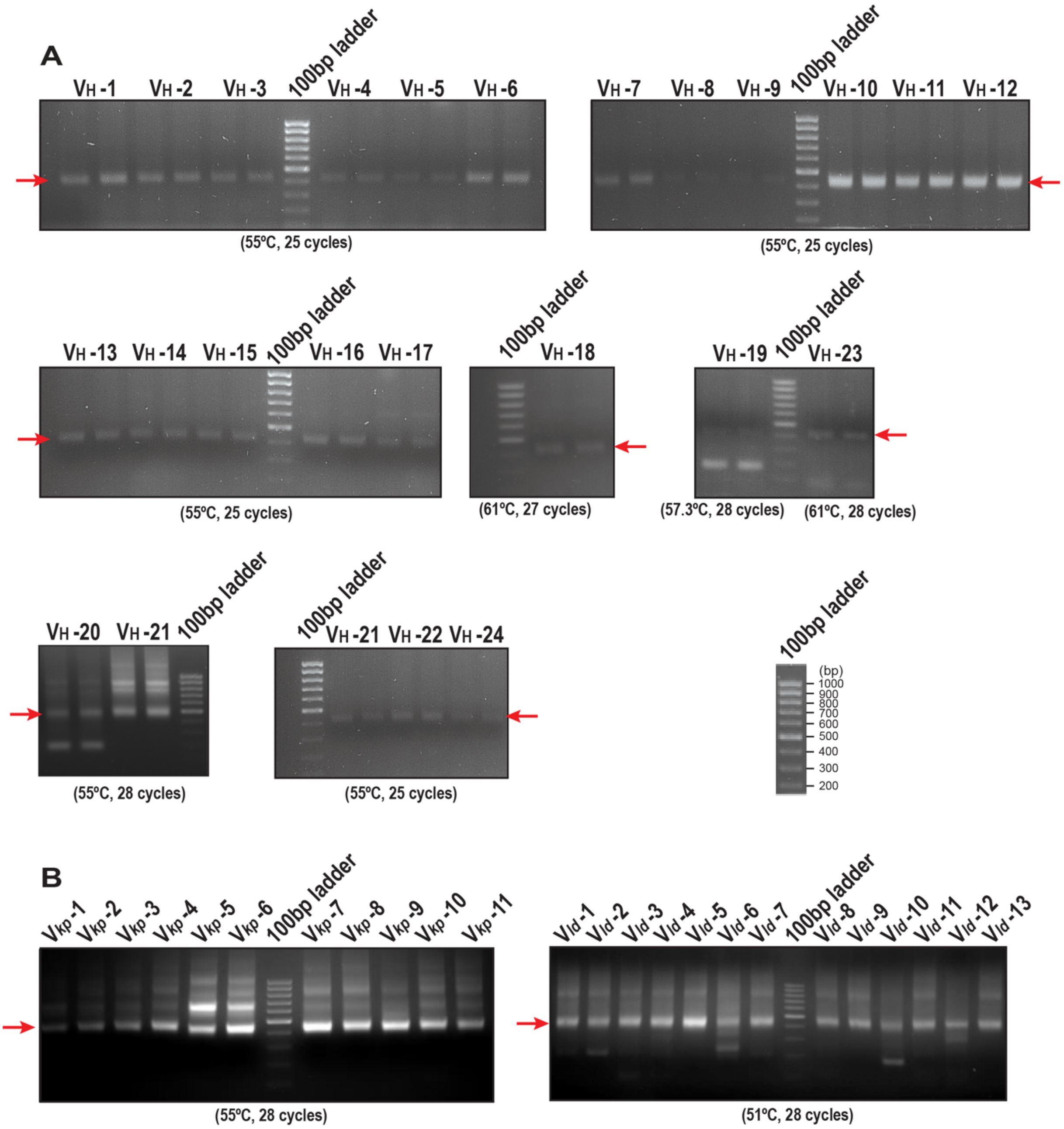

**Figure S2**. PCR products of VH and VL genes. Electrophoresis agarose gels illustrating the VH genes (A) and VL(B). These fragments were amplified from cDNA derived from lymphocytes of 50 healthy donors and 25 AML patients. The observed bands are ~400 bp in size. 100pb DNA ladder (Thermofisher) was used. Each band signifies 100 bp, with 1000 bp for the uppermost band and 100 bp for the lowest band.

**Supplemental Figure S3**

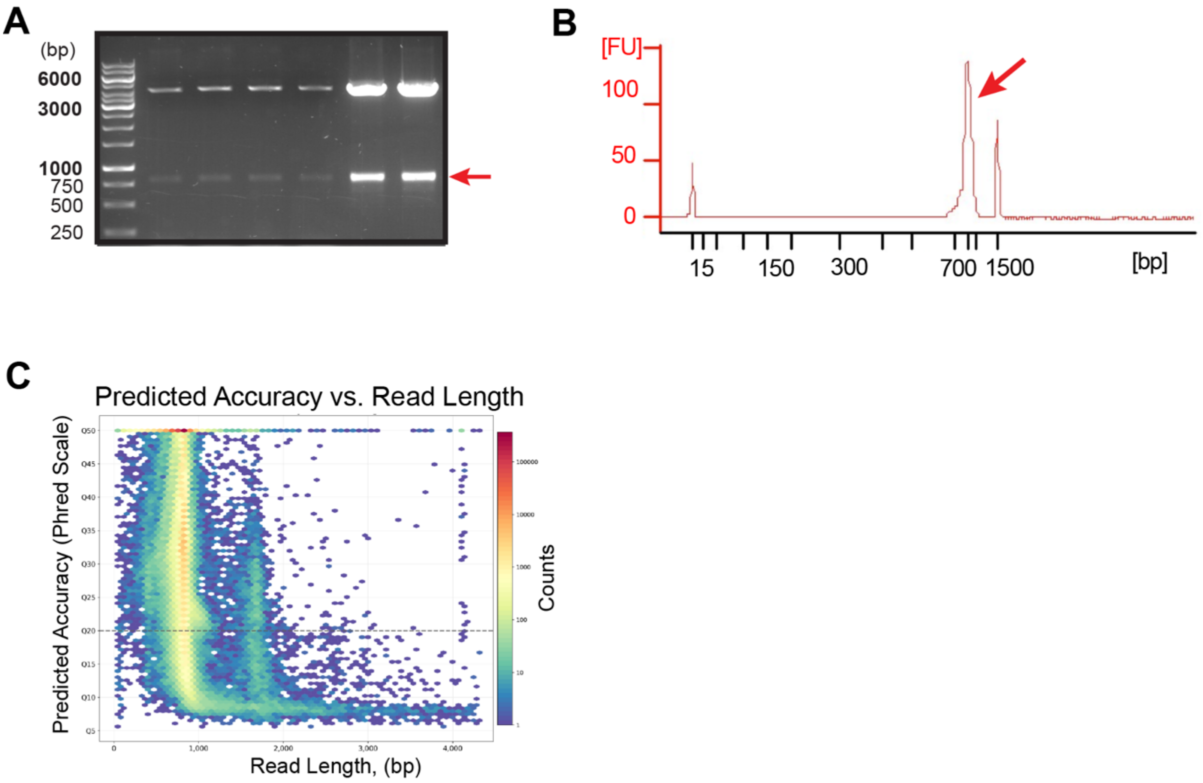

**Figure S3**. High-throughput sequencing of the scFv phagemids library. (A) Agarose gel electrophoresis displaying the excised scFv bands from the vector for PacBio sequencing, with an approximate size of ~800 bp. (B) Bioanalyzer analysis of the purified scFv samples submitted for PacBio sequencing. The peak at ~800 bp, pointed out with the red arrow, corresponds to the expected size for scFv fragments. The left and right peaks represent the used 15 bp and 1500 bp ladders, respectively. (C) Predicted Accuracy vs Read Length. The boundary between HiFi Reads and other CCS Reads is shown as a dashed line at QV 20.

### Supplemental Figure S4

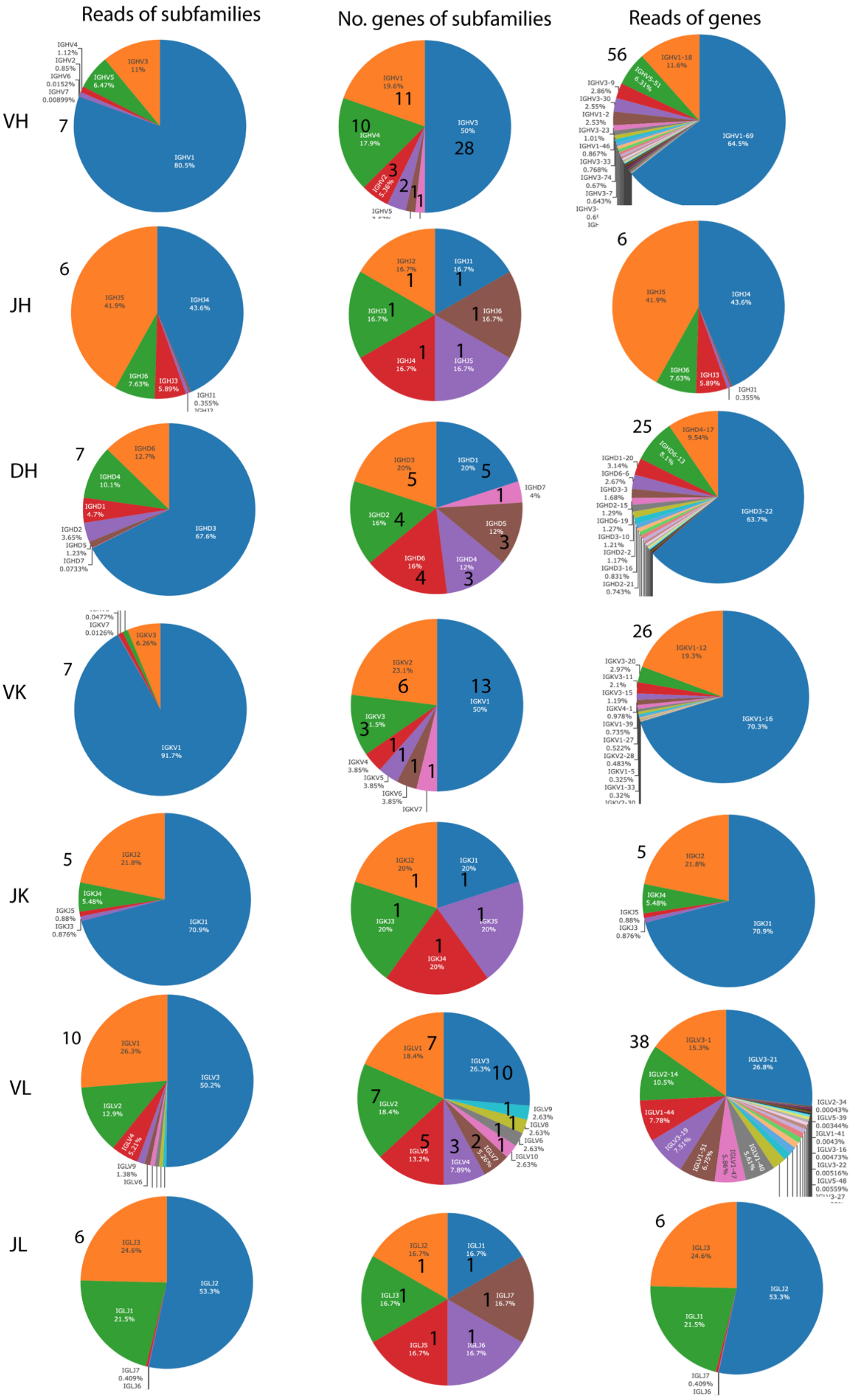

| | HV | HD | HJ | KV | KJ | LV | LJ |
|---|---|---|---|---|---|---|---|
| No.genes | 56 | 25 | 6 | 26 | 5 | 38 | 6 |
| No.subfamilies | 7 | 7 | 6 | 7 | 5 | 10 | 6 |

**Figure S4**. Frequency and distribution of subfamilies and genes of heavy chains and light chains from the PacBio sequencing. VH: variable genes/subfamilies of heavy chains; DH: diversity genes/subfamilies of heavy chains; JH: joining genes/subfamilies of heavy chains; VK: variable genes/subfamilies of kappa light chains; JK: joining genes/subfamilies of kappa light chains; VL: variable genes/subfamilies of lambda light chains; JK: joining genes/subfamilies of lambda light chains;

**Supplemental Figure S5**

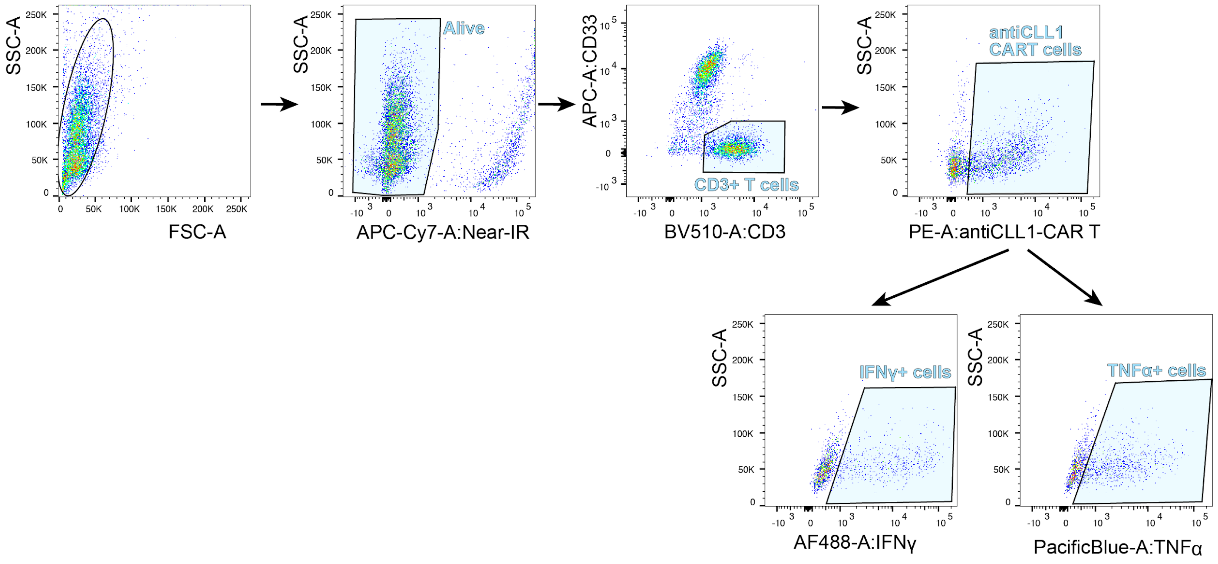

**Figure S5.** ICS assay gating approach. Prior to analysis, CAR T cells were cocultured with target cells for 6 hours. Here shows an example of anti-CLL-1-CAR T carrying scFv 6 (C6) cocultured with MV4-11 cells. Antibodies used for ICS assay are as labeled in the image.

**Supplemental Figure S6**

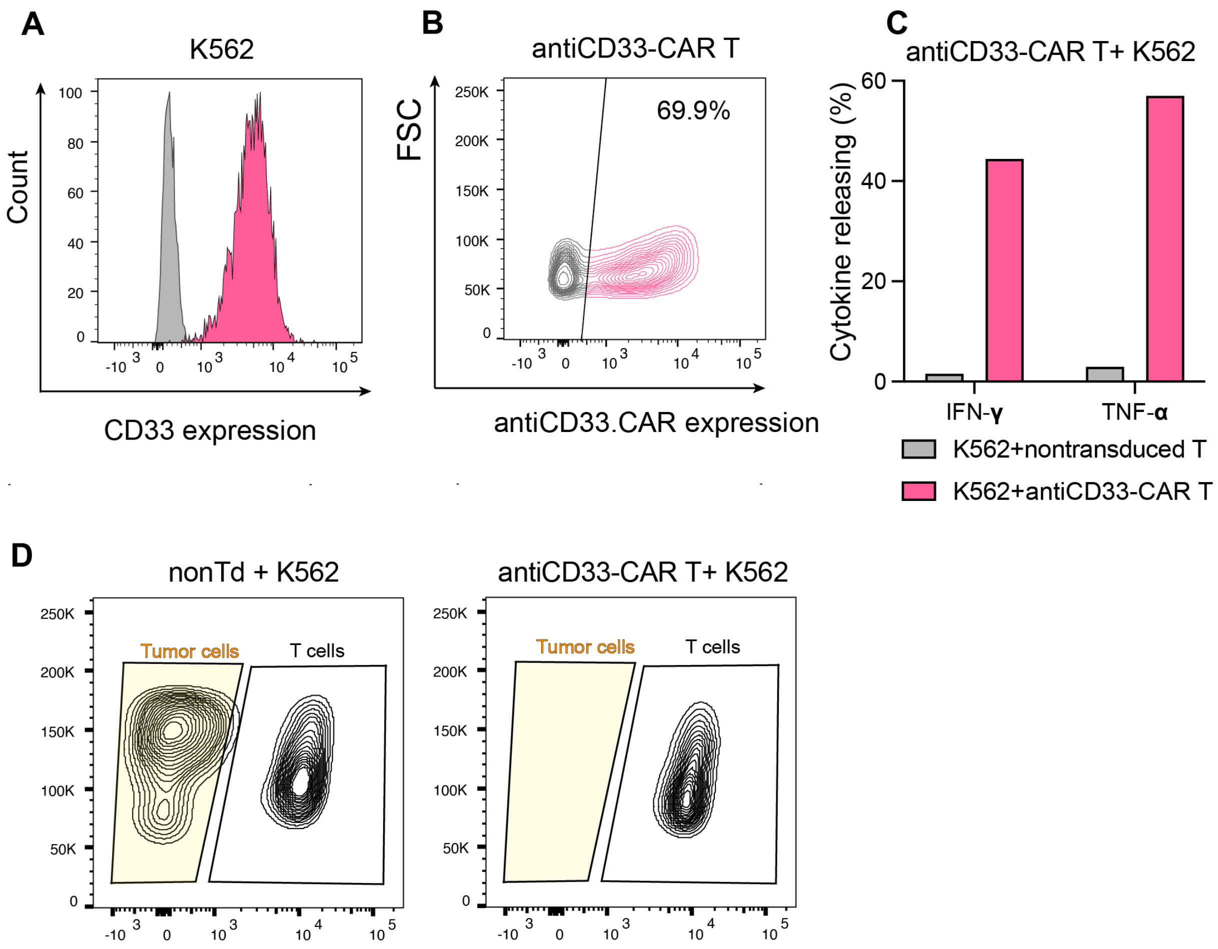

**Figure S6.** K562 cells are antiCD33.CAR T cells sensitive. (A) Flow cytometry (FC) analysis of CD33 expression on K562 cells. Prior to FC, cells were stained with APC-antiCD33 antibody (clone 67.6) (magenta) for 15 min at room temperature in dark. Unstained control is shown in grey. (B) FC analysis of antiCD33-CAR expression (magenta) on transduced T cells. Cells were stained with PE-antiCAR. (C) Intracellular cytokine releasing of antiCD33-CAR T cells stimulated by K562 cells. (D) Short-time killing of antiCD33-CAR T cells against K562 cells. Cells were measured 2 days after coculture

### Supplemental Table S1. Primers used for scFv-phagemid library generation

| | | |
|---|---|---|
| $V_H$-1 | NcoI-VH1a-F | gtcctcgca cc atg g cc CAG GTG CAG CTG GTG CAG TCT GG |
| $V_H$-2 | NcoI-VH1b-F | gtcctcgca cc atg g cc CAG GTT CAG CTG GTG CAG TCT GG |
| $V_H$-3 | NcoI-VH1c-F | gtcctcgca cc atg g cc CAG GTC CAG CTT GTG CAG TCT GG |
| $V_H$-4 | NcoI-VH1d-F | gtcctcgca cc atg g cc CAG GTC CAG CTG GTA CAG TCT GG |
| $V_H$-5 | NcoI-VH1e-F | gtcctcgca cc atg g cc GAG GTC CAG CTG GTA CAG TCT GG |
| $V_H$-6 | NcoI-VH1f-F | gtcctcgca cc atg g cc CAA ATG CAG CTG GTG CAG TCT GG |
| $V_H$-7 | NcoI-VH1g-F | gtcctcgca cc atg g cc CAG ATG CAG CTG GTG CAG TCT GG |
| $V_H$-8 | NcoI-VH2a-F | gtcctcgca cc atg g cc CAG ATC ACC TTG AAG GAG TCT GG |
| $V_H$-9 | NcoI-VH2b-F | gtcctcgca cc atg g cc CAG GTC ACC TTG AAG GAG TCT GG |
| $V_H$-10 | NcoI-VH3a-F | gtcctcgca cc atg g cc GAG GTG CAG CTG GTG GAG TCT GG |
| $V_H$-11 | NcoI-VH3b-F | gtcctcgca cc atg g cc GAA GTG CAG CTG GTG GAG TCT GG |
| $V_H$-12 | NcoI-VH3c-F | gtcctcgca cc atg g cc CAG GTG CAG CTG GTG GAG TCT GG |
| $V_H$-13 | NcoI-VH3d-F | gtcctcgca cc atg g cc GAG GTG CAG CTG TTG GAG TCT GG |
| $V_H$-14 | NcoI-VH3e-F | gtcctcgca cc atg g cc GAG GTG CAG CTG GTG GAG ACT GG |
| $V_H$-15 | NcoI-VH3f-F | gtcctcgca cc atg g cc GAG GTG CAG CTG GTG GAG TCC GG |
| $V_H$-16 | NcoI-VH3g-F | gtcctcgca cc atg g cc GAG GTG CAG CTG GTG GAG TCT CG |
| $V_H$-17 | NcoI-VH4a-F | gtcctcgca cc atg g cc CAG GTG CAG CTG CAG GAG TCG GG |
| $V_H$-18 | NcoI-VH4b-F | gtcctcgca cc atg g cc CAG CTG CAG CTG CAG GAG TCC GG |
| $V_H$-19 | NcoI-VH4c-F | gtcctcgca cc atg g cc CAG CTG CAG CTA CAG CAG TGC GG |
| $V_H$-20 | NcoI-VH4d-F | gtcctcgca cc atg g cc CAG CTG CAG CTA CAG CAG TCG GG |
| $V_H$-21 | NcoI-VH5a-F | gtcctcgca cc atg g cc GAA GTG CAG CTG GTG CAG TCT GG |
| $V_H$-22 | NcoI-VH5b-F | gtcctcgca cc atg g cc GAG GTG CAG CTG GTG CAG TCT GG |
| $V_H$-23 | NcoI-VH6-F | gtcctcgca cc atg g cc CAG GTA CAG CTG CAG CAG TCA GG |
| $V_H$-24 | NcoI-VH7-F | gtcctcgca cc atg g cc CAG GTG CAG CTG GTG CAA TCT GG |
| $V_H$-Rev | HindIII-VH-R | gtcctcgca aag ctt GAC CGA TGG GCC CTT GGT GGA |
| V*kp*-1 | MluI-V*k*1-F1 | accgcctcc a cgc gta RAC ATC CAG ATG ACC CAG TCT CC |
| V*kp*-2 | MluI- V*k*1-F2 | accgcctcc a cgc gta GMC ATC CRG WTG ACC CAG TCT CC |
| V*kp*-3 | MluI-V*k*1-F3 | accgcctcc a cgc gta GTC ATC TGG ATG ACC CAG TCT CC |
| V*kp*-4 | MluI-V*k*2-F1 | accgcctcc a cgc gta GAT ATT GTG ATG ACC CAG ACT CC |
| V*kp*-5 | MluI-V*k*2-F2 | accgcctcc a cgc gta GAT RTT GTG ATG ACT CAG TCT CC |
| V*kp*-6 | MluI-V*k*3-F1 | accgcctcc a cgc gta GAA ATW GTG WTG ACR CAG TCT CC |
| V*kp*-7 | MluI-V*k*3-F2 | accgcctcc a cgc gta GAA ATT GTA ATG ACA CAG TCT CC |
| V*kp*-8 | MluI-V*k*4-F | accgcctcc a cgc gta GAC ATC GTG ATG ACC CAG TCT CC |
| V*kp*-9 | MluI-V*k*5-F | accgcctcc a cgc gta GAA ACG ACA CTC ACG CAG TCT CC |
| V*kp*-10 | MluI-V*k*6-F1 | accgcctcc a cgc gta GAA ATT GTG CTG ACT CAG TCT CC |
| V*kp*-11 | MluI-V*k*6-F2 | accgcctcc a cgc gta GAT GTT GTG ATG ACA CAG TCT CC |
| *kp*Rev | NotI-V*kappa*-R | accgcctcc gc ggc cgc gaa gac aga tgg tgc agc cac agt |

| | | |
|---|---|---|
| V*l*d-1 | MluI-V*l*1-F1 | accgcctcc a cgc gta CAG TCT GTG CTG ACT CAG CCA CC |
| V*l*d-2 | MluI-V*l*1-F2 | accgcctcc a cgc gta CAG TCT GTC YTG ACG CAG CCG CC |
| V*l*d-3 | MluI-V*l*2-F | accgcctcc a cgc gta CAG TCT GCC CTG ACT CAG CCT |
| V*l*d-4 | MluI-V*l*3-F1 | accgcctcc a cgc gta TCC TAT GWG CTG ACW CAG CYA C |
| V*l*d-5 | MluI-V*l*3-F2 | accgcctcc a cgc gta TCT TCT GAG CTG ACT CAG GAC CC |
| V*l*d-6 | MluI-V*l*3-F3 | accgcctcc a cgc gta TCC TAT GAG CTG ATG CAG CCA CC |
| V*l*d-7 | MluI-V*l*4-F1 | accgcctcc a cgc gta CTG CCT GTG CTG ACT CAG CCC |
| V*l*d-8 | MluI-V*l*4-F2 | accgcctcc a cgc gta CAG CYT GTG CTG ACT CAA TCR YC |
| V*l*d-9 | MluI-V*l*5-F | accgcctcc a cgc gta CAG SCT GTG CTG ACT CAG CC |
| V*l*d-10 | MluI-V*l*6-F | accgcctcc a cgc gta AAT TTT ATG CTG ACT CAG CCC CA |
| V*l*d-11 | MluI-V*l*7/8-F | accgcctcc a cgc gta CAG RCT GTG GTG ACY CAG CAG CC |
| V*l*d-12 | MluI-V*l*9-F | accgcctcc a cgc gta CAG CCT GTG CTG ACT CAG CCA CC |
| V*l*d-13 | MluI-V*l*10-F | accgcctcc a cgc gta CAG GCA GGG CTG ACT CAG CCA CC |
| V*l*d-Rev | NotI-V*lambda*-R | Accgcctcc gc ggc cgc aga gga sgg ygg gaa cag agt gac |

Nucleotide W: A/T; Y: C/T; R: G/A; S: G/C; M: A/C

### Supplemental Table S2. Subreads statistics of PacBio sequencing

Each polymerase read is partitioned to form one or more subreads, which contains sequence from a single pass of a polymerase on a single strand of an insert within a SMRTbell™ template and no adapter sequences. The subreads contain the full set of quality values and kinetic measurements

| Sample | SMRT Cell name | Bases (bp) | Total Bases (G) | Reads Number | Mean Length (bp) | Longest (bp) | N50 (bp) |
|---|---|---|---|---|---|---|---|
| ABL2022 | m64164_220406_095920 | 164158643767 | 164.16 | 267789854 | 613 | 577571 | 856 |

**Note:**

**Sample:** Sample name.
**Cell:** SMRT Cells name.
**Bases(bp):** The base number.
**Total Bases(G):** The base number of all cell.
**Reads Number:** The number of reads.
**Mean Length(bp):** The average length of all reads.
**Longest(bp):** The longest read length of all reads.
**N50(bp):** 50% of all reads are longer than the value.

### Supplemental Table S3. CCS reads statistics of PacBio sequencing

Circular Consensus Sequencing (CCS) Analysis computes consensus sequences from multiple "passes" around a circularized single DNA molecule (SMRTbell® template). CCS Analysis uses the Arrow framework to achieve optimal consensus results given the number of passes available.

| Sample | SMRT Cell name | Bases (bp) | Total Bases (G) | Reads Number | Mean Length (bp) | Longest (bp) | N50 (bp) |
|---|---|---|---|---|---|---|---|
| ABL2022 | m64164_220406_095920 | 10413946303 | 10.41 | 1538864 | 6767 | 539126 | 64837 |

**Note:**

**Sample:** Sample name.
**Cell:** SMRT Cells name.
**Bases(bp):** The base number.
**Total Bases(G):** The base number of all cell.
**Reads Number:** The number of reads.
**Mean Length(bp):** The average length of all reads.
**Longest(bp):** The longest read length of all reads.
**N50(bp):** 50% of all reads are longer than the value.

**Supplemental Table S4. HiFi reads statistics of PacBio sequencing**

PacBio highly accurate long reads, known as HiFi reads, are produced by sequencing a single molecule of DNA multiple times. HiFi reads can be used across a wide range of SMRT Sequencing applications, from whole genome sequencing for de novo assembly, comprehensive variant detection, RNA sequencing and more. HiFi reads are generated with CCS Analysis whose quality value is equal to or greater than 20.

| Sample | SMRT Cell name | Bases (bp) | Total Bases (G) | Reads Number | Mean Length (bp) | Longest (bp) | N50 (bp) | Median Quality | Passes Number |
|---|---|---|---|---|---|---|---|---|---|
| ABL2022 | m64164_220406_095920 | 558586831 | 0.56 | 693364 | 805 | 16287 | 823 | Q60 | 24 |

**Note:**

**Sample:** Sample name.
**Cell:** SMRT Cells name.
**Bases(bp):** The base number.
**Total Bases(G):** The base number of all cell.
**Reads Number:** The number of reads.
**Mean Length(bp):** The average length of all reads.
**Longest(bp):** The longest read length of all reads.
**N50(bp):** 50% of all reads are longer than the value.
**Median Quality:** The median number of CCS Reads whose quality value is equal to or greater than 20.
**Passes Number:** The mean number of passes used to generate CCS Reads whose quality value is equal to or greater than 20
